# Drug delivery to rodents: how to deal with body mass and water intake fluctuations?

**DOI:** 10.1101/615567

**Authors:** Gonçalves Leidyanne Ferreira, Fernandes-Santos Caroline

## Abstract

**Introduction:** Animal models are used to test the safety and efficacy of drugs. They are often administered to rodents in the drinking water, but it has some limitations, such as the drug stability, variations of water consumption and body mass. We investigated telmisartan (TEL) stability in mice drinking water by UV spectrophotometry, and if water intake and body mass fluctuations change drug ingestion.

**Methods:** Female C57BL/6 mice at two months old, were fed for eight weeks with a purified AIN93M diet, or a high-fat high-sucrose diet (HFHS). TEL 5 mg/Kg/day was administered ad libitum to mice in the drinking water during three weeks concomitant with diets, summing 11 weeks of diet feeding.

**Results:** UV spectrophotometry could detect TEL at the wavelength of 300 nm, and it remained stable in mice drinking water for seven days, at the concentration expected. Mice gain weight after eight weeks on high-fat high-sucrose diet feeding, and TEL 5 mg/kg/day in the drinking water for three weeks reduced it. TEL did not change water intake. Not adjusting TEL concentration weekly would lead to a higher intake of TEL by mice.

**Discussion:** We demonstrated that body mass and water intake fluctuations significantly change the amount of drug that the animal receive, and it would add a bias to the experiment. TEL remains stable for at least seven days in wrapped mice water bottles in the animal care facility, and UV spectrophotometry proved to be a simple and low-cost method to detect TEL in mice drinking water.

## INTRODUCTION

Animal models are widely used to understand the onset and development of diseases. They are also used to test the safety and efficacy of drugs, as well as the pathophysiology and molecular mechanisms underlying drug effect on body systems. Drugs can be delivered to rodents in many ways, such as diluted in the drinking water and chow, into the stomach by oral gavage, and intraperitoneally. Each route has its advantages and disadvantages, and the most commonly used routes are oral gavage [1, 2] and drinking water [3-5]. Drinking water is the easiest route for drug delivery since it does not stress the animal, it does not require specific skills or training, and it has low cost and low risk for the animal compared to other methods. On the other hand, rodents might not receive the planned drug dose, because water intake varies along the experiment. Also, in obesity studies where rodents gain or lose weight, drug dosage is also affected. Another concern is drug stability since some substances lost their activity when exposed to light. Thus, the researcher must assure that the animal ingests the planned dose and that the drug solution is stable.

Telmisartan (TEL) is an antihypertensive drug approved by the US Food and Drug Administration (FDA) since 1998 [6]. It is an angiotensin II type 1 (AT1) receptor blocker and it is chemically described as 4’-[(1,4’dimethyl-2’-propyl [2,6’-bi-1H-benzimidazol]-1’yl) methyl]-[1,1’-biphenyl]-2-carboxylic acid [7, 8]. TEL is prescribed to reduce blood pressure and to prevent cardiovascular morbidity and mortality since it reduces left ventricular hypertrophy, arterial stiffness and atrial fibrillation [8, 9]. TEL also exerts pleiotropic effects on glucose metabolism and insulin sensibility due to its action as a partial peroxisome proliferator activated receptor γ (PPARγ) agonist [1, 8]. Many studies have been conducted in rodent models of hypertension [10, 11], diabetes [12, 13] and obesity [14, 15] to investigate how TEL acts on blood pressure and glucose metabolism. TEL reduces body mass in rodents, depending on the dose and how long it is offered. Thus it is important to assure that the animals ingested the planned dose. Another concern is that TEL is light sensitive.

There are many methods to analyze TEL concentration in solution, such as ultraviolet (UV) spectrophotometry, immunoassay, liquid chromatography-mass spectrometry (LC-MS) [16], high performance liquid chromatography (HPLC) [17, 18], ultraperformance liquid chromatography (UPLC) [19], and also polarography and visible spectrophotometry [16]. Except for spectrophotometry, these techniques are expensive, requires high-cost equipment, demands expert training, and uses solvents that can be harmful to health if incorrectly used [16]. Thus, the British Pharmacopoeia and the Indian Pharmacopoeia recommends UV spectrophotometry for TEL analysis [19]. Therefore, our goal was to exploit how to assure the planned drug dosage when the drug is administered to mice that are subjected to water intake and body weight fluctuations throughout the experiment, using TEL as an example. Also, we evaluated if it is possible to measure TEL concentration and stability in the drinking water offered to mice. To accomplish that, we used the method of UV spectrophotometry proposed by Chavhan to measure TEL concentration in mice drinking water [20].

## MATERIALS AND METHODS

### Ethical aspects

The local Ethics Committee approved the handling and experimental protocols to Care and Use of Laboratory Animals (CEUA#647/15). The study was performed following the Animal Research Reporting In vivo Experiments ARRIVE guidelines and the Guideline for the Care and Use of Laboratory Animals (US NIH Publication N° 85-23. Revised 1996) [21].

### Animal husbandry and diet

Twenty female C57BL/6 mice at two months old were maintained in collective polycarbonate microisolators of 30×20×23/28×18×11 cm external/internal dimensions, with a wire bar lid that serves as food hopper and water bottle holder on ventilated racks (Scienlabor Industrial Equipment, SP, Brazil). Five mice were housed per cage. The housing conditions were 12 h light/dark cycles, 21±2°C, 60±10% humidity and air exhaustion cycle of 15 min/h. At three months old, mice were feed for eight weeks with a purified AIN93M diet [22], or a high-fat high-sucrose diet (HFHS) modified from the AIN93M diet (Pragsolucoes, Jau, Sao Paulo, Brazil) to induce weight gain. TEL 5 mg/Kg/day was administered ad libitum to mice in the drinking water during three weeks concomitant with diets, summing 11 weeks of diet feeding (Fig. 1).

**Figure 1.**
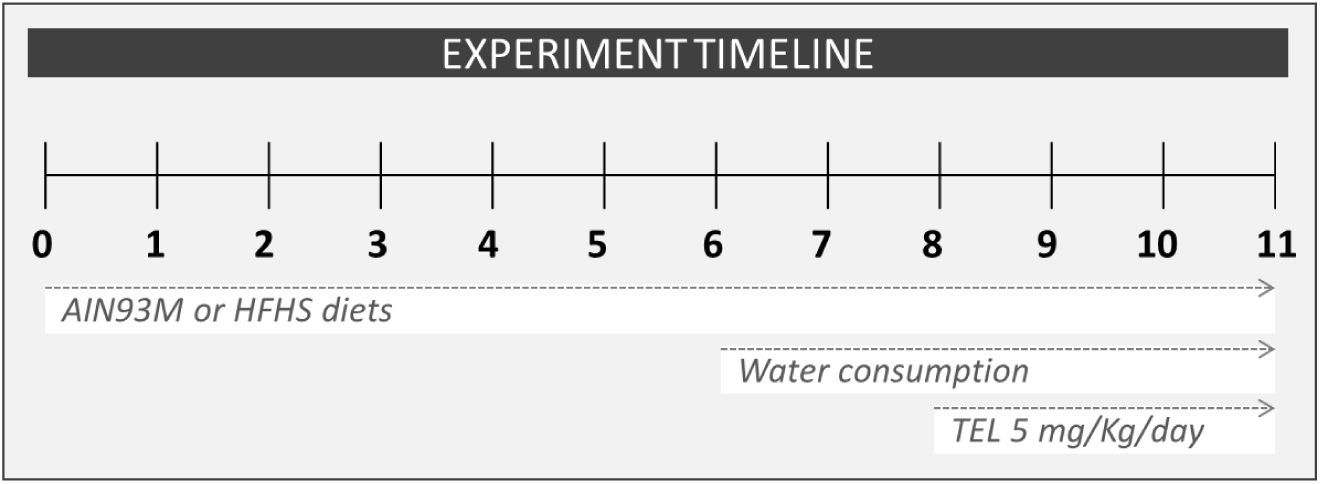
Experiment timeline (weeks). Mice were fed during 11 weeks with control AIN93M diet or a modified AIN93M diet rich in fat and sucrose (HFHS). Water intake was measured along two weeks before TEL administration, and then twice weekly during TEL administration. TEL was offered for three weeks in the drinking water at 5 mg/Kg/day. Abbreviations: HFHS, high-fat high-sucrose diet; TEL, telmisartan.

### Instrumentation

The spectrophotometric measurements were carried out using an Epoch UV-visible light spectrophotometer (BioTek^®^ Instruments Inc, Vermont, USA). It consists of a photodiode detector with a xenon flash light source. Optical performance is λ range of 200-999 nm, ± 2 nm accuracy, ± 0.2 nm repeatability, 5 nm bandpass, 0.000 to 4.000 optical density (OD). Readings were performed in 96-well. An analytical scale (Shimadzu, AUW220D, Kyoto, Japan) was used for reagent weighing.

### Reagents

TEL 80 mg (Lot.2680246, Ranbaxy Laboratories Limited, Mohali, Punjab, India) was bought in a local market. Milli-Q purified water was used for reagent dilution (Millipore, Massachusetts, EUA).

### Solution preparation for tel assay

Solution preparation was based on Chavhan *et al.* 2013. They showed that TEL has better solubility in 0.1N NaOH. Also, TEL (Ranbaxy Laboratories Limited tablet) concentration claimed in the label matched the estimated concentration of 100 mg/tablet by 99.15% after six determinations [20]. Thus, we used TEL tablets to prepare the standard and working solutions for TEL determination in drinking water.

#### Standard stock solution (1 000 µg/mL)

Average weight of 5 tablets was determined. The powder equivalent to 50 mg of TEL was weighted and dissolved in 25 mL 0.1N NaOH by sonication (570 seconds, 42 KHz, 60 watts, Ultrasonic Cleaner CD-4800, Practical Systems Inc, Florida, USA). Then, it was filtered through filter paper of 14 µm pore, and volume was made up to 50 mL with 0.1N NaOH to give a 1 000 µg/mL stock solution.

#### Working stock solution (100 µg/mL)

2.5 mL was withdrawn from stock standard solution and diluted to 25 mL with 0.1N NaOH to get a 100 µg/mL solution.

#### Selection of analytical wavelength

The absorbance of working solution and blank was sampled in the range of UV radiation from 200-400 nm at 5 nm intervals. TEL showed maximum absorption at the wavelength (λ_max_) 300 nm (Fig. 2 A).

**Figure 2.**
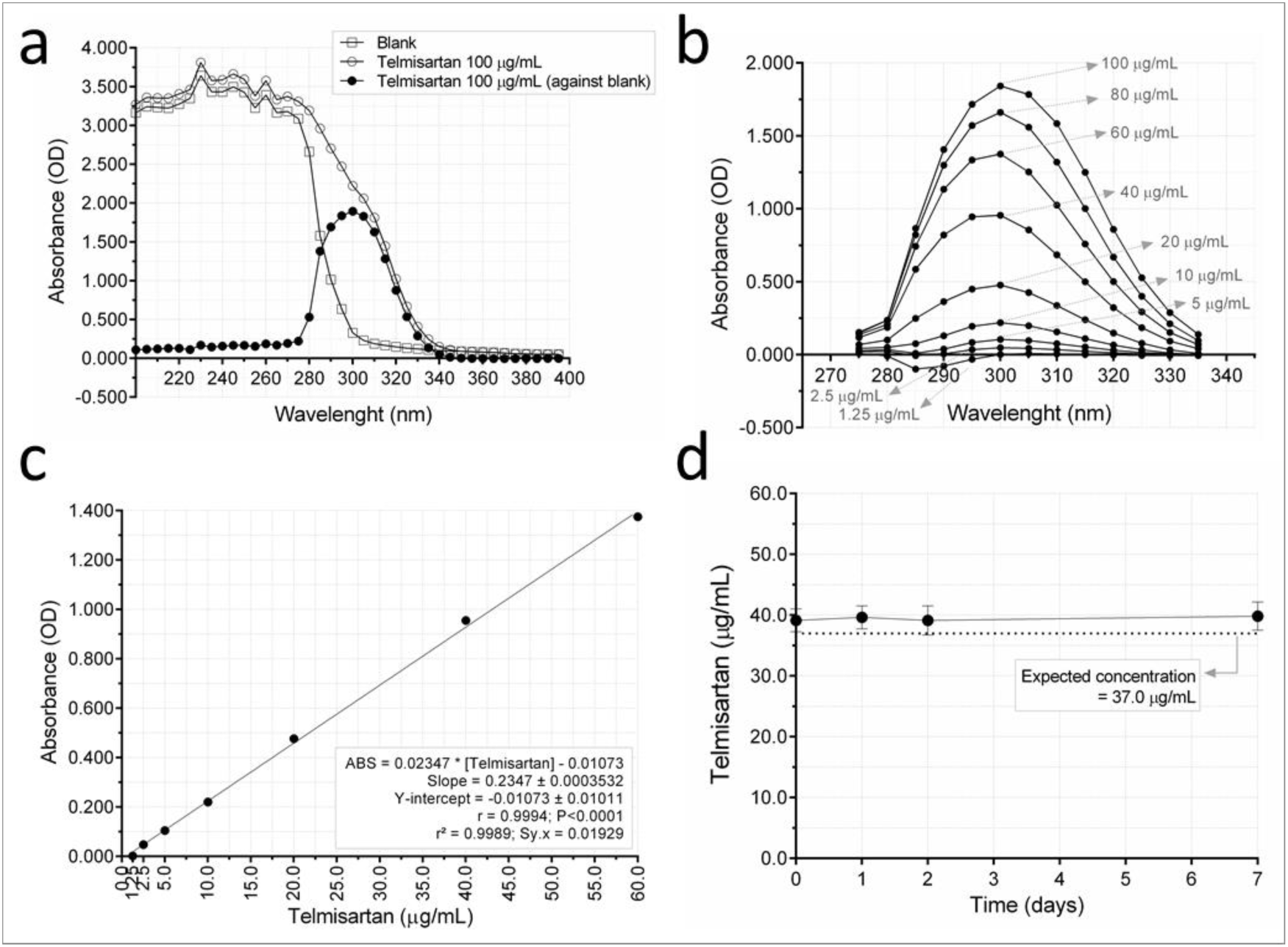
TEL detection by UV spectrophotometry. **A**, TEL showed maximum absorption at 300 nm. **B**, Absorption of TEL solution in the UV spectrum (200-400 nm) of different concentrations (1.25-100 µg/mL). **C**, At 300 nm, TEL showed linear response with increasing concentration. **D**, Observed TEL concentration (mean±SD) of four mice bottles that were left in the animal care facility for seven days wrapped in aluminum foil (one sample t-test).

#### Calibration curve (1.25-100 µg/mL)

Appropriate dilutions of TEL working solution were made to obtain a concentration range of 1.25, 2.5, 5, 10, 20, 40, 60, 80, 100 µg/mL in Milli-Q purified water. In a 96-well microplate, 300 µL of each solution was read in the range of 200-400 nm at 5 nm intervals in triplicates, and the absorbance at 300 nm was used for standard curve calculation (Fig. 2 B). Calibration solutions showed linear response with increasing concentration. Curve range of 1.25 to 60 µg/mL was chosen, based on the goodness of fit of the linear regression (R squared=0.9989, Sy.x 0.01929) and Pearson correlation (r=0.9994, P<0.0001) (Fig. 2 C).

### TEL dilution in mice drinking water

Previous literature shows that drinking water is the better route for TEL administration to mice [3, 5, 23]. Due to space limitation, mice are often housed in collective cages. Therefore, water intake is evaluated per cage, and water consumption represents the average of five mice, not individual consumption. Water consumption was measured daily from Monday to Friday throughout two weeks before TEL administration to estimate average water consumption. Daily water consumption was used to estimate the amount of TEL required to prepare the TEL drinking solution. We needed to assure that the volume of water ingested by each mouse in one day had the amount of TEL planned (5 mg/Kg/day). Water intake was also monitored during the experiment since changes in water consumption would change TEL dosage.

Table 1 shows how the amount of TEL powder (macerated tablet) required to be diluted in mice drinking water was determined. Firstly, it was calculated how much TEL (drug itself) would be necessary daily for mice (110.0 µg/day/mice), based on average body weight of mice housed in the same collective cage (22.0 g). Secondly, the result was corrected based on the amount of pure TEL presented in the TEL powder (in this example, 660.0 µg/day/mice of TEL powder). Finally, to know the amount of TEL powder required for dilution in mice drinking water, it was necessary to know daily water intake (3.2 mL/day/mice, average of five mice housed in a same cage) to assure that the planned TEL quantity (that it, 666.0 µg of TEL powder) is presented in 3.2 mL of water.

**Table 1.** Simulation showing how to calculate Telmisartan weight required to prepare mice drinking water

| **TELMISARTAN DOSE CALCULATION \*** |  |
| --- | --- |
| Body weight | 22.0 g |
| Telmisartan dose | 5.0 mg/kg/day |
| Telmisartan (tablet powder) | 80 mg in 479.8 mg powder |
| Telmisartan (pure) | 110.0 µg/day/mice |
| Telmisartan (powder) | 660.0 µg/day/mice |
| **DRINKING WATER PREPARATION \*** |  |
| Water volume | 300.0 mL/bottle |
| Water consumption | 3.2 mL/day/mice |
| Telmisartan required (powder) | 61.85 mg |

\* Body weight and water consumption are illustrative to understand calculations, based on averaged data obtained during experiment.

TEL was offered to mice in water bottles wrapped in aluminum foil to protect the solution from exposure to environmental light since TEL is light sensitive. TEL was offered ad libitum in 300 mL water bottles for three weeks to evaluate its impact on body mass. Body mass was weighted weekly. The amount of TEL diluted in mice drinking water was adjusted weekly based on the average body mass and water consumption of the previous week for each mice cage.

### TEL concentration and stability in mice drinking water

Since our concern about TEL concentration and stability emerged after the experiment was finished, we designed an assay to simulate the conditions to which TEL solution was subjected during experimentation. Thus, four mice water bottles were filled with a TEL solution consisting of 22.15 mg of TEL tablet powder diluted in 100 mL of filtered water (∼36.93 µg/mL of pure TEL).

Water bottles were wrapped in aluminum foil and left at animal’s room facility for one week. One sample of each bottle was collected at 0, 1, 2, and 7 days after solution preparation to evaluate TEL concentration and stability. In a 96-well microplate, 300 µL of filtered water (blank), TEL solution or TEL calibration curve solutions (1.25-60 µL/mL) were read at 300 nm in triplicates. TEL concentration was calculated from the standard curve at 300 nm.

### Expected and observed TEL concentration in mice drinking water

To precisely offer the intended (expected) amount of TEL requires the knowledge of individual body mass and water intake. Since mice were housed in collective cages, we knew their body mass, but not water intake because the last is an average per cage. It might be a critical issue for drug delivery because some mice would consume more (or less) drug compared to what was planned by the investigator. Thus, we first evaluated if using the average water intake per cage would significantly change the intended (expected) TEL dosage for each mouse. For this analysis, we calculated the expected TEL concentration using individual mouse body mass and average water intake per cage, and the observed TEL concentration was calculated based on average body mass and water intake per cage. Secondly, we investigated if the observed TEL concentration would be much different from expected if water intake and body mass were not corrected weekly. For this analysis, the expected TEL concentration was also calculated using individual mouse body mass and average water intake per cage, whereas the observed TEL concentration was based on average water intake and body mass per cage measured before TEL treatment started, and it was not corrected along the next three weeks.

### Statistical analysis

Data are expressed as mean ± SD and analyzed for normality and homoscedasticity of variances. Linear regression was used to generate the standard curve and to calculate TEL concentration. One-way ANOVA of Kruskal-Wallis and post hoc Dunn’s multiple comparisons was used to compare water consumption among cages. Water consumption and body mass were analyzed by repeated measures ANOVA with post hoc Sidak’s multiple comparison test and linear trend. Observed and expected TEL concentration was compared by one sample t-test, paired t-test and Bland-Altman plot. A *P*-value <0.05 was considered statistically significant (GraphPad^®^ Prism software v. 6.0, La Jolla, CA, USA).

## RESULTS

### TEL detection by uv spectrophotometry and stability in mice drinking water

We confirmed that TEL is detectable by UV spectrophotometry. The working stock solution showed maximum absorption at the wavelength (λmax) of 300 nm (Fig. 2A). This wavelength was used to determine the optical density of a series of TEL dilutions (Fig. 2B) to establish the standard curve (Fig. 2C). Once the standard curve was established, TEL concentration in four water bottles containing a known concentration of TEL (37.0 µg/mL) was assessed, to evaluate if it would remain stable along seven days (Fig. 2D). The observed and expected TEL concentrations were similar, showing that TEL remains stable in mice drinking water for at least seven days.

### TEL did not change water intake and induced body weight loss

Water intake was assessed two weeks before TEL offering to mice and also during the three weeks of TEL intake, to evaluate if TEL would change water consumption (Table 2). Despite tight control of animal care facility conditions, water intake usually displays small fluctuations among weeks, and for this reason, we solely compared average water intake among cages on the same week. Water intake was similar among cages, showing that TEL does not interfere with it. Eight weeks on HFHS diet feeding lead to body mass gain, and three weeks of TEL administration reduced it (Fig. 3A). In summary, we found that water intake was lightly increased and body mass decreased in the last three weeks of the experiment. For this reason, TEL concentration in mice drinking water was intentionally reduced, otherwise mice would receive more TEL than the initial dose planned of 5 mg/Kg/day.

**Table 2.** Average water intake (mL/mice) before and after Telmisartan administration to mice

| Week | Control Tel |  | HFHS Tel |  | P |
| --- | --- | --- | --- | --- | --- |
|  | Cage 1 | Cage 2 | Cage 3 | Cage 4 |  |
| -2 | 3.3±0.7 (20.6) | 3.1±0.5 (15.6) | 3.9±1.0 (23.7) | 3.9±1.0 (24.2) | 0.53 |
| -1 | 3.2±0.6 (19.1) | 3.3±0.4 (10.9) | 3.8±0.4 (10.1) | 3.7±0.4 (9.5) | 0.51 |
| 0 | 3.6±0.7 (20.3) | 3.8±0.8 (21.1) | 4.1±1.2 (29.5) | 4.1±1.0 (25.2) | 0.92 |
| 1 | 4.0±0.5 (12.9) | 4.0±0.5 (12.6) | 4.0±0.5 (12.1) | 4.3±0.4 (12.1) | 0.54 |
| 2 | 4.0±0.4 (10.2) | 4.5±0.4 (8.5) | 4.5±0.7 (16.1) | 4.7±0.7 (15.7) | 0.26 |
| 3 | 4.9±1.2 (23.7) | 4.1±0.4 (9.7) | 4.1±0.2 (5.8) | 4.6±0.4 (7.9) | 0.50 |

Five mice were allocated per cage. Average water intake represents daily measurements from Monday to Friday (total of 5 readings), except for the last week, where cages 1 and 2 were read three days and cages 3-4 read 4 days.
Data is expressed as mean±SD (C.V.). Non-parametric one-way ANOVA Kruskal-Wallis and post hoc Dunn's multiple comparison tests were performed. Comparison was not made between weeks since it is expected that water intake presents time-dependent fluctuation.

**Figure 3.**
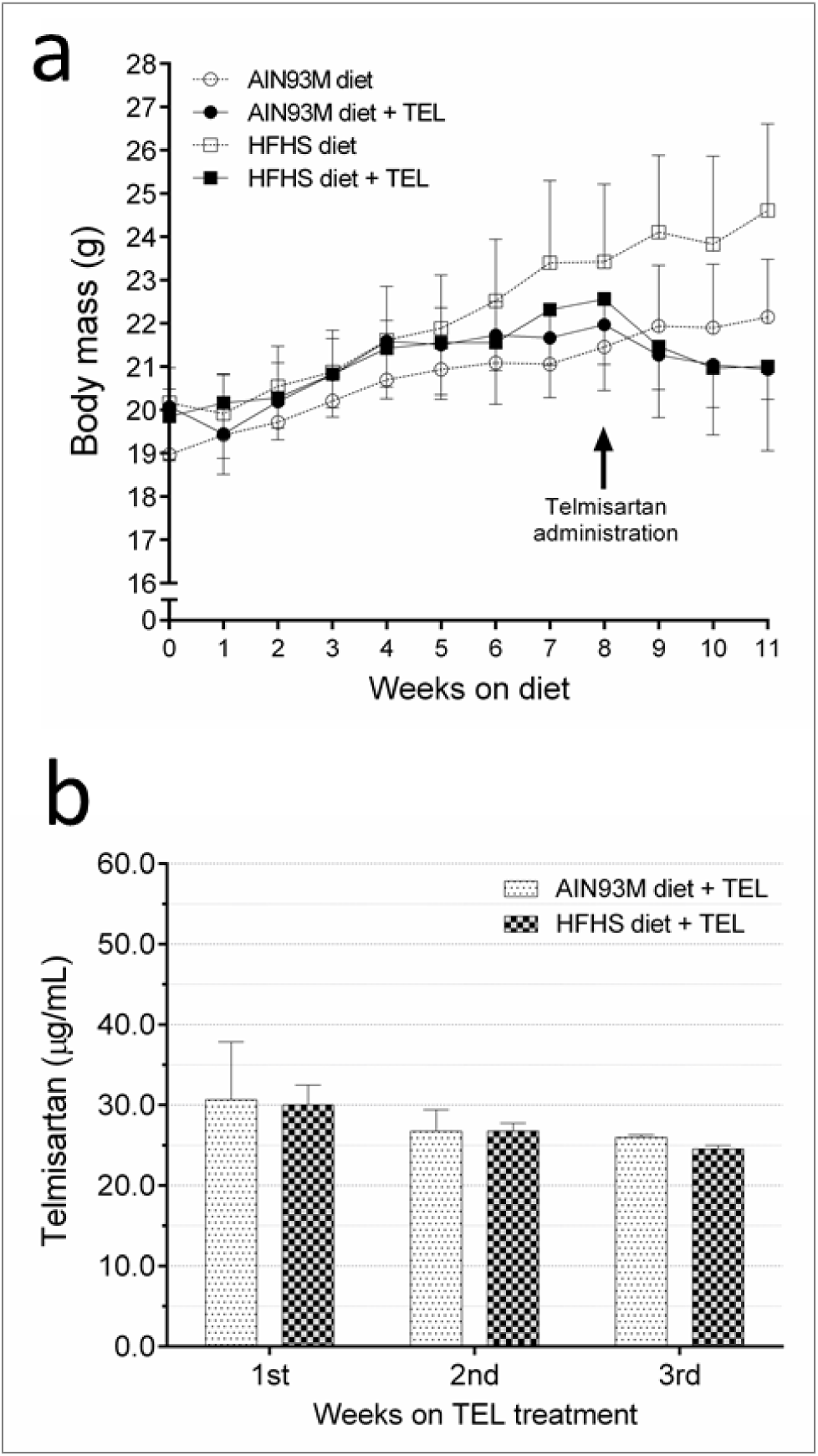
Body mass and TEL dose adjustment. **A**, Body mass of mice fed for 11 weeks with control AIN93M diet or a modified AIN93M diet rich in fat and sucrose (HFHS). From 8^th^ to 11^th^ week, TEL was offered to mice in the drinking water. [*] indicates HFHS diet vs. HFHS diet +Tel, P<0.05, t-Student test. **B**, Adjustment of TEL concentration in mice drinking water due to TEL-induced body weight loss. Statistical comparison was not performed since n=2 cages/group, because mice were housed in collective cages.

### Observed and expected TEL concentration in mice drinking water

Figure 4 shows the expected (planned) and observed TEL concentration in mice drinking water in two different situations. In the first scenario, TEL concentration was adjusted every week based on mouse body mass (individual) and water intake (cage average), and as a result, observed and expected TEL concentrations were similar (Fig. 4A). In the second scenario (Fig. 4B), we plotted the observed TEL concentration in mice drinking water if we had used only the values of body mass and water intake before treatment started. Therefore, not considering the temporal fluctuations in mice body mass and water intake, resulted in a significant difference among observed and expected TEL concentration in mice drinking water of the HFHS diet +TEL group. In this second approach, observed TEL concentration would be 13% higher (P=0.0059) in the second week of experiment and 16% higher (P=0.002) in the third week. Mice fed with the AIN93M diet showed no significant difference in observed and expected TEL concentration, and it is likely because TEL did not lead to significant changes in body mass for this group. Comparing these two scenarios (adjusted versus non-adjusted TEL concentration) using the Bland-Altman plot, we noticed that not adjusting TEL concentration weekly (Fig. 4D) increased the bias and the difference between expected and observed TEL concentration, compared to when it is done (Fig. 4C).

**Figure 4.**
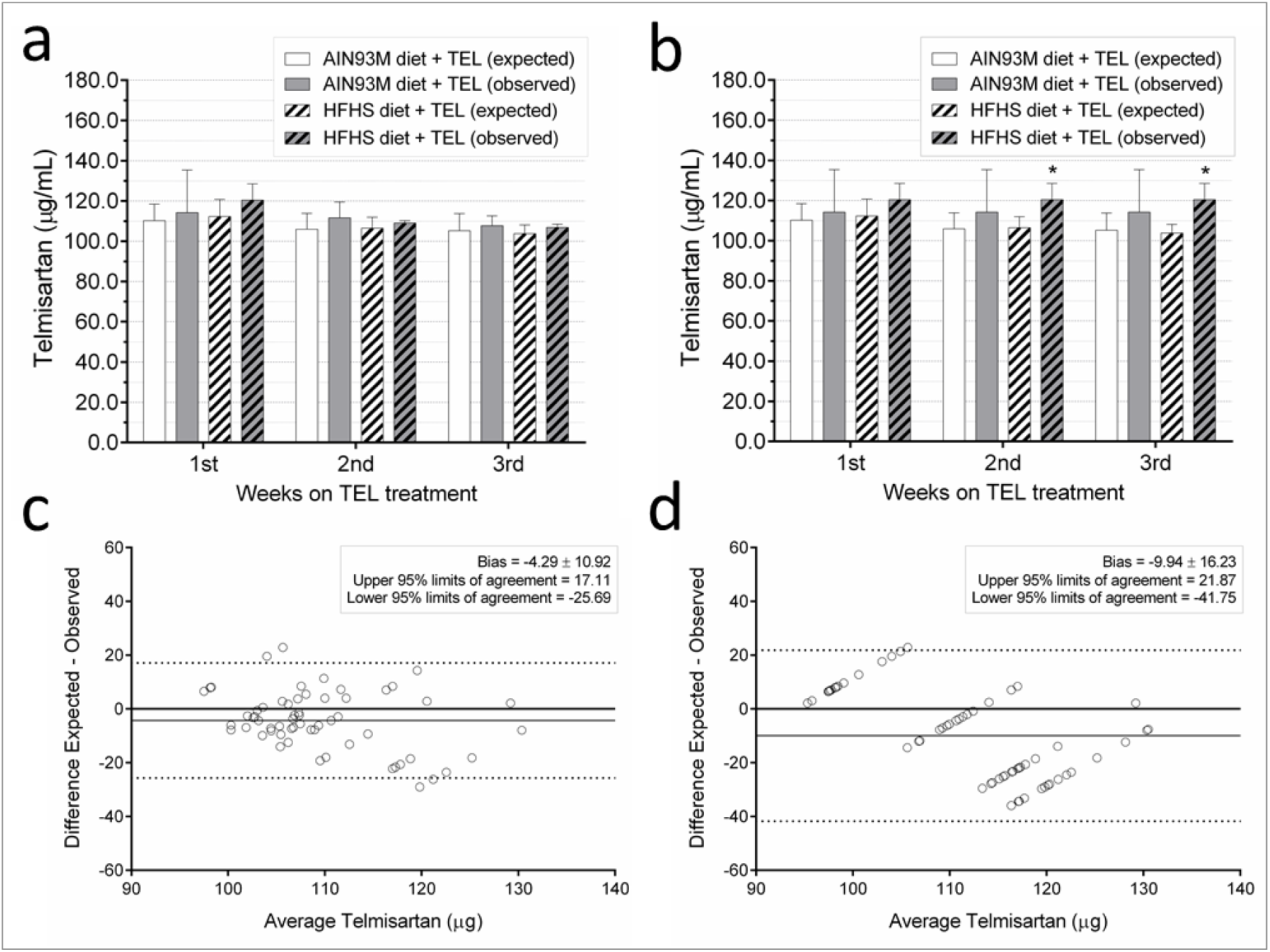
Expected versus observed TEL concentration in the drinking water offered to female C57BL/6 mice. **A**, Average TEL concentration in mice drinking water was calculated twice weekly. **B**, Average TEL concentration in mice drinking water, but it was based on average pre-treatment water intake and body mass thus it was not calculated twice weekly along with the experiment. **C-D**, Bland-Altman plot among observed and expected TEL concentration in mice drinking water. In C, TEL concentration was recalculated twice weekly, but not on D. When indicated, P<0.05, comparing expected vs. observed data (paired t teste). Abbreviations: AIN93M, control diet; HFHS, high-fat high-sucrose diet, TEL, telmisartan.

## DISCUSSION

We investigated how to guarantee drug dosage when the compound is administered to mice in drinking water, knowing that body mass and water intake might change throughout the experiment, and used TEL as an example. We saw that UV spectrophotometry could detect TEL at a wavelength of 300 nm and it is stable in mice water bottles wrapped with aluminum foil for at least seven days. If body mass and water intake changes throughout the experiment, it significantly changes the amount of drug that the animals will receive, adding a bias to the experiment.

In the last decades, researchers have been developing and validating methods to quantify TEL in bulk and pharmaceutical dosage forms. Patel *et al.* estimated TEL dosage by UV spectrophotometry using methanol as solvent [19]. They showed that TEL has maximum absorption at wavelength 296 nm, but in 233 nm it is also possible to detect a peak, and linearity was observed between 4-16 μg/mL. Pandey *et al.* used TEL diluted in 0.1N NaOH and found a maximum absorption at 234 nm and linearity between 4-24 μg/mL [24]. Chavhan *et al.* also used TEL diluted in 0.1N NaOH and found a maximum absorption at 295 nm and linearity between 2-12 μg/mL [20]. In our study, we followed the protocol described by Chavhan *et al.* and have found a maximum absorption at 300 nm. Regarding TEL linearity range for detection, we decided for 1.25-60 μg/mL based on the goodness of curve fit of the linear regression. Our range is larger compared to the previously published data, but the authors mentioned above did not mention if they tested TEL concentrations below or above the ranges they reported on their works.

Regarding stability, Patel *et al.* showed high precision for TEL identification along three consecutive days [19]. Our data corroborate with it since TEL concentration remained stable along seven days in mice drinking water. As TEL is light sensitive, wrapping the bottle with aluminum foil likely assured its stability, as well as the controlled environmental temperature of the animal care facility (21±2°C). Patel *et al.* investigated TEL degradation under diverse conditions to estimate its stability (acidic, alkali, neutral, oxidation, photolytic, and thermal conditions), and they found that TEL undergoes degradation in all the stressing conditions tested. Its stability was evaluated in a neutral solution (double distilled water) for 5 hours, and authors found 10% degradation. However, the solution was placed in the oven at 70°C for this assay [19].

It’s interesting to note that, in a posterior review by Patel *et al*., it was discussed that TEL determination by UV method could suffer interference due to other UV absorbing compounds. Therefore, the HPLC and HPTLC methods would be better indicated for this kind of analysis. However, they are expensive and require elaborate procedures, which can make it difficult to implement [25]. Thereby, the UV spectrophotometry method is an easy, simple and economical way to quantify TEL to evaluate its concentration in mice drinking water. We believe that the presence of other compounds might be a challenge for TEL determination in blood serum for instance, but not when using water, NaOH or methanol as solvents.

The most common route for TEL administration is oral by ad libitum intake in the drinking water [3, 5, 23]. TEL dosage to C57BL/6 mice was chosen based on both Human equivalent dose and after an extensive review of several articles. First, we considered the Human dose of 40 mg/day, which is equivalent to approximately 8.2 mg/kg/day for mice [26]. The dose range used in the literature ranges from 3-10 mg/Kg/day. Based on this two information, we have chosen an intermediate dose of 5 mg/Kg/day, which has been used in other studies with C57BL/6 mice and have shown biological effects on preventing body weight gain and body fat accumulation [4, 23].

In Humans, TEL average terminal elimination half-life is between 20-24 hours, contributing to the 24 hours antihypertensive efficacy with once-daily dosing [7-9, 27]. In mice, the terminal elimination half-life is 8-10 hours [7], which might be critical when it is provided in the drinking water since animals would constantly be exposed to the compound. It seems that rodents on pelleted diet consume most of their water immediately before and after they eat food [28]. When housed under standard 12-h light/12-h dark cycle, mice consume most of their food during the dark, with short bouts during the light cycle [29]. To bias elimination, an alternative would be to administer TEL twice or three times a day by oral gavage. However, it is necessary that the investigator be trained on the technique, since unskilled personnel might stress and hurt the animal, by traumatic injury related to unappropriated restrain, by damaging the oral cavity, esophagus or trachea, or the solution might get access to the trachea and thus the lungs, leading to animal death.

Zhang *et al.* described a protocol for voluntary oral administration of drugs to mice, where the drug is offered in artificially sweetened and flavored jelly and given to mice that have been trained to eat the jelly [30]. They report that mice need to be trained for 2-4 days, and they need to be individually housed to assure that every mouse eat the jelly and they do not fight for it, especially for male mice. The jelly is made of a mixture of Splenda^®^ low caloric sweetener, water, gelatin and flavoring essence imitation (strawberry of chocolate). This method is of interest since each jelly can be prepared with a different drug concentration based on mouse body weight, what is not possible for drugs delivered in the drinking water. On the other hand, it is time-consuming, considering that one experimental group receiving the drug would have 8-10 mice that will be likely treated for at least one to several weeks.

Before introducing a drug into the drinking water, the investigator must determine the strain’s average 24-hour water consumption. Ad libitum water intake must be determined for the same strain, sex, age, and weight of the rodent strain that the investigator is willing to work [31, 32]. In our experience, male C57BL/6 mice with 3-6 months old drink about 4-8 mL/day of water, and females about 3-5 mL/day. If they are submitted to a high-salt diet, water intake increases significantly, ranging from 8-20 mL/day for male and 7-14 for female C57BL/6 mice (authors unpublished data and [33]). Thus, water intake needs to be continuously monitored throughout the experiment, to guarantee that the planned drug dose in indeed received. We measured water consumption two weeks before the beginning of the experiment, but for the simplest determination, one may use three consecutive days [31]. To evaluated water intake, rodents must be housed in their usual cage to keep their usual routine. The water bottle that animals are already accustomed is filled and weighted using a digital scale with 0.1g accuracy. The bottle is weighed three days in a row in the same time of the day to determine the 24-hour ad libitum water consumption; the measures are averaged and then divided by the number of rodents in the cage.

We did not notice expressive changes in water intake among the four mice cages. Corroborating with our data, TEL offered ad libitum to male Sprague-Dawley rats into their drinking water did not change water intake [3]. Depending on the drug used, it can have good palatability for mice, and thus water consumption is increased. The opposite is also true, and therefore the planned dose will vary. Additionally, it is important to monitor food consumption. Fluctuations in food intake might change water intake, and also the compound itself might lead to an increase or decrease in food consumption. A decrease (or increase) in food consumption may change body mass and be a confounding factor to data interpretation regarding drug impact on body mass and body composition. Another example is the use of fructose solution to induce insulin resistance and hypertension in rodents [34]. In our experience, fructose offered to the rodent in drinking water increases water consumption. Therefore, care must be taken if the investigator is willing to administer a drug such as TEL into the drinking water when there is another factor influencing water intake (e.g., high-salt diet or fructose) because drug consumption will also change from what was planned.

As exposed, fluctuations in water intake and body mass have an impact on drug consumption. Therefore drug dose needs to be adjusted weekly. We showed that TEL decreased the body mass of mice from HFHS diet + TEL group. If TEL concentration were not adjusted, mice would be exposed to a higher dose than expected, if we consider that water intake did not change. Our study evaluated three weeks of TEL treatment, but in longer experiments, if mice display a persistent weight loss, TEL dosage would be consistently increased and thus enhancing its effects on body systems, adding a huge bias to data interpretation. The lack of TEL concentration adjustment did not impact mice receiving the AIN93M diet + TEL, and it is likely because they did not display an important variation in body mass, compared to the HFHS diet + TEL group. Thus, if body mass did not change throughout drug intervention, drug adjustment is not a critical issue.

Since TEL is offered ad libitum, it would be interesting to determine its blood concentration. In mice, peak plasma concentration (C_max_) after one dose of 1 mg/Kg of TEL is 162 ng/mL after oral administration [7], but to date, we did not find a report for its plasma concentration when administered ad libitum to mice. Regarding methods to detect TEL, Virkar *et al.* developed and validated an HPLC method with fluorescence detection in rabbit plasma [35]. Salama *et al.* also reported the use of HPLC-UV in Human plasma, and was able to detect between 1-10 µg/mL of TEL [36]. Finally, an LC-MS method was developed for TEL determination in Human plasma [37], but it requires a large blood sample, which is a limitation for mice studies. Overall, these techniques are expensive for laboratories that do not use it as routine. We attempted to determine TEL in mice serum using UV spectrophotometry, but unsuccessfully, since it required large blood samples to establish a concentration curve based on mice serum as a matrix, and also pure TEL instead of powder from tablet maceration.

## CONCLUSIONS

In conclusion, we demonstrated that body mass and water intake fluctuations throughout the experiment significantly change the amount of drug that the animal receive, and it would add a bias to the experiment. TEL remains stable for at least seven days in wrapped mice water bottles in the animal care facility, and UV spectrophotometry proved to be a simple and low-cost method to detect TEL in mice drinking water.

## ACKNOWLEDGMENTS

Authors are thankful to Rye Watanabe, Luana Tavares Acioli and Larissa Guedes Rodrigues for their help in animal care.

## AUTHORS’ CONTRIBUTIONS

L.F.G performed the experiment, collected, analyzed, and interpreted data, and wrote the manuscript draft. C.F-S designed the experiment and revised the manuscript. All authors read and approved the final manuscript.

## LIST OF ABBREVIATIONS

AT1: angiotensin II type 1 receptor
C: control diet
C-Tel: control diet + Treatment
FDA: US Food and Drug Administration
HFHS: high-fat high-sucrose diet
HFHS-Tel: HFHS diet + Treatment
HPLC: high-performance liquid chromatography
LC-MS: liquid chromatography-mass spectrometry
PPARγ: peroxisome proliferator-activated receptor γ
TEL: telmisartan
UV: ultraviolet
UPLC: ultraperformance liquid chromatography

